# Reference mouse strain assemblies for BALB/c Nude and NOD/SCID mouse models

**DOI:** 10.1101/2023.03.16.532783

**Authors:** Emanuel Schmid-Siegert, Mengting Qin, Huan Tian, Bulak Arpat, Bonnie Chen, Ioannis Xenarios

**Affiliations:** NGS-AI Switzerland; NGS-AI Suzhou

## Abstract

Mouse xenograft models play a vital role in tumor studies for research as well as for screening of drugs for the pharmaceutical industry. In particular, models with compromised immunity are favorable to increase the success of transplantation, such as e.g. NOD/SCID and BALB/c Nude strains. The genomic sequence and alterations of many of these models still remain elusive and might hamper a model’s further optimization or proper adapted usage. This can be in respect to treatments (e.g. NOD/SCID sensitivity to radiation), experiments or analysis of derived sequencing data of such models. Here we present the genome assemblies for the NOD/SCID and BALB/c nude strains to overcome this short-coming for the future and improve our understanding of these models in the process. We highlight as well first insights into observed genomic differences for these models compared to the C57BL/6 reference genome. Genome assemblies for both are close to full chromosome representations and provided with liftover annotations from the GRCm39 reference genome.

## Introduction

Reference genomes have been essential in many research fields and with the advances in high throughput sequencing, detailed characterizations of genomes become more readily available [***Miga et al., 2020***], including the mouse genome [***Gregory et al., 2002***]. The description and understanding of the genomic structure of mice have been extended recently by ***Ferraj et al. [2022***] in-depth characterization of 20 genetically distinct inbred mice strains. Their goal was to deepen the understanding of causative alleles for phenotypic variation across individuals.

One field where understanding the biology, and therefore genomic makeup, of the model strain remains key, is in the generation of immunocompromised mice. These mouse strains play a crucial role in establishing mouse tumor models such as cell line-derived xenografts (CDXs), patient derived-xenografts (PDXs) and homografts in syngeneic mice [***Olson et al., 2018, Okada et al., 2019, Li et al., 2017***]. The genetic background of a strain profoundly affects the phenotype and potential applications of a mouse model, which contributes to defects in innate immunity such as reduced natural killer (NK) cell activity, absence of circulating complement, and deficits in macrophages as well as antigen-presenting cells [***Chuprin et al., 2023***].

The combination of such defects are often obtained through crosses of single defect models such as in the case of NOD and SCID crossed to a NOD/SCID mouse model. Some of the defects are genomically well defined and derive for example from a single point mutation, others from transgenes or less well characterized genomic alterations. Subsequently, the genomic sequence and alterations of some of these models are more elusive and might hamper it’s further genetic and immunonolgic optimization. Based on PDX-recycled data, we present here the genomic characterization of the BALB/c Nude and NOD/SCID models and their reference genomes to facilitate use and understanding of these models. The mouse strains used typically for PDX models at Crown Biosciences are BALB/c Nude and NOD/SCID, for which the genome has till today not been characterized to the full extend. Non-obese diabetic Severe combined immunodeficiency (NOD/SCID, ***Bosma et al. [1983***])-based immunocompromised mice lack both, functional T- and B-lymphocytes and present a dysfunctional natural killer cell (NK) activity. BALB/c Nude mice on the other hand have lower macrophage-mediated phagocytosis of human cells and a deficiency of mature T-lymphocytes, which makes them together with the nude phenotype suitable for transplantation.

Sequencing data from multiple PDXs was pooled and the mouse strain assembled after eliminating human sequences. As tissue derived from PDX samples is always a mixture of host and graft material, this is reflected similarly in the obtained sequencing data. Depending on multiple factors (location, type of tumor, experimentator and more) this ratio has been seen to be as low as < 1% but as well high as beyond 60 % [***Karamboulas et al., 2020, Woo et al., 2019***]. The latter is less desired for sequencing as it lowers the targeted assay coverage and complicates analysis of the data. Recent progress has been made to evaluate this already prior sequencing by using a high-throughput, low-cost, NGS-based screening approach [***Chen et al., 2020***]. This approach has not been applied to the models used in this study.

We present a strategy to make use of the unwanted host-derived data to reconstruct it’s corresponding genome *de novo*. Particular care has to be taken to eliminate residual human contamination at different steps for raw data and assembled genomic data. Applying that strategy we present close to full-chromosome assemblies for the BALB/c Nude and NOD/SCID strains as a resource to improve our understanding of their genomic make-up and facilitate their use in xenograft related studies.

## Results

### Sample characterization

Due to their length combined with high quality, we chose PacBio HIFI reads for the genome assembly process. This feature combination reduces significantly the possibility to accidentally introduce human sequences into the assembly step compared to short reads or more error-prone long reads. The HIFI reads were screened with xengsort, a fast and accurate xenograft sorting algorithm [***Zentgraf and Rahmann, 2021***], and reads which were unambiguously identified as graft (human) were removed. All the other reads were kept for further analysis, which included as well the categories “both” and “ambiguous”. The former including reads which carry unique kmers for both organism whereas the latter containing kmers which are equally present in both organism - e.g. telomeric-, centromeric- or very conserved regions. This resulted in a 19 % to 62% observed mouse-content per sample (supplementary table S1) in agreement with previous reports [***Chen et al., 2021, Rossello et al., 2013***]. Median mouse-content were consistent across BALB/c Nude (24.5%) and NOD/SCID (31%) samples.

We confirmed the genotype of each PDX sample based on known marker mutations. The BALB/c Nude mouse model is characterized by a 1bp deletion in the exon 3 of the Foxn1 gene which leads to a truncated protein and subsequently the nude phenotype [***Schlake, 2001***]. The NOD/SCID mice should not carry the nude mutation but another marker mutation which is in the 2nd to last exon of the Prkdc gene and results in an early stop codon of that gene [***Blunt et al., 1996***]. This NOD/SCID mutation is not present in the BALB/c Nude model. We systematically analyzed these 2 positions for all 13 samples using two approaches based on the mapping of HIFI reads to the reference GRCm39 genome: i) *de novo* SNV calling on the entire genome and ii) pileup analysis at the marker positions without any genotype inference. The expected genotype was confirmed in all samples by both approaches.

Prior to pooling samples for the assembly process, we investigated further their similarity and variation for both, within- and in-between-strains. To this end, we called single nucleotide variants (SNV) and structural variants (SV) from the alignment of the HIFI reads against the GRCm39 reference genome using deepvariant [***Poplin et al., 2018***] and PacBio’s pbsv algorithm [***Wenger et al., 2019***], respectively. Variants in tandem repeated regions in the reference genome were removed, as well as in regions of problematic alignments. To further reduce noise due to shallow depth, variants with low confidence scores were excluded and only homozygous germline variants were further analyzed. Comparison of number of shared events (stranded) between samples identified a clear separation between BALB/c Nude and NOD/SCID mice using SVs (supplementary figure S1) and SNVs (supplementary figure S2). Within strain, samples were very similar to each other, underlining the low sample heterogenity on the genomic makeup. Interestingly, less variant of each type were overall detected in BALB/c Nude mice compared to NOD/SCID mice.

### Global variant characterization

To further analyze and understand the genomic variations, we grouped homozygous SV- and SNV-events into three groups: events observed i) commonly in both strains, ii) uniquely in BALB/c Nude and iii) uniquely in NOD/SCID (Figure 1). As expected, no events were reported on the Y chromosome which is absent in our uniquely female sample collection. Similar to SNVs, SV events were more present in the NOD/SCID mice compared to the BALB/c Nude ones, most prominent for deletion and translocation events.

**Figure 1.**
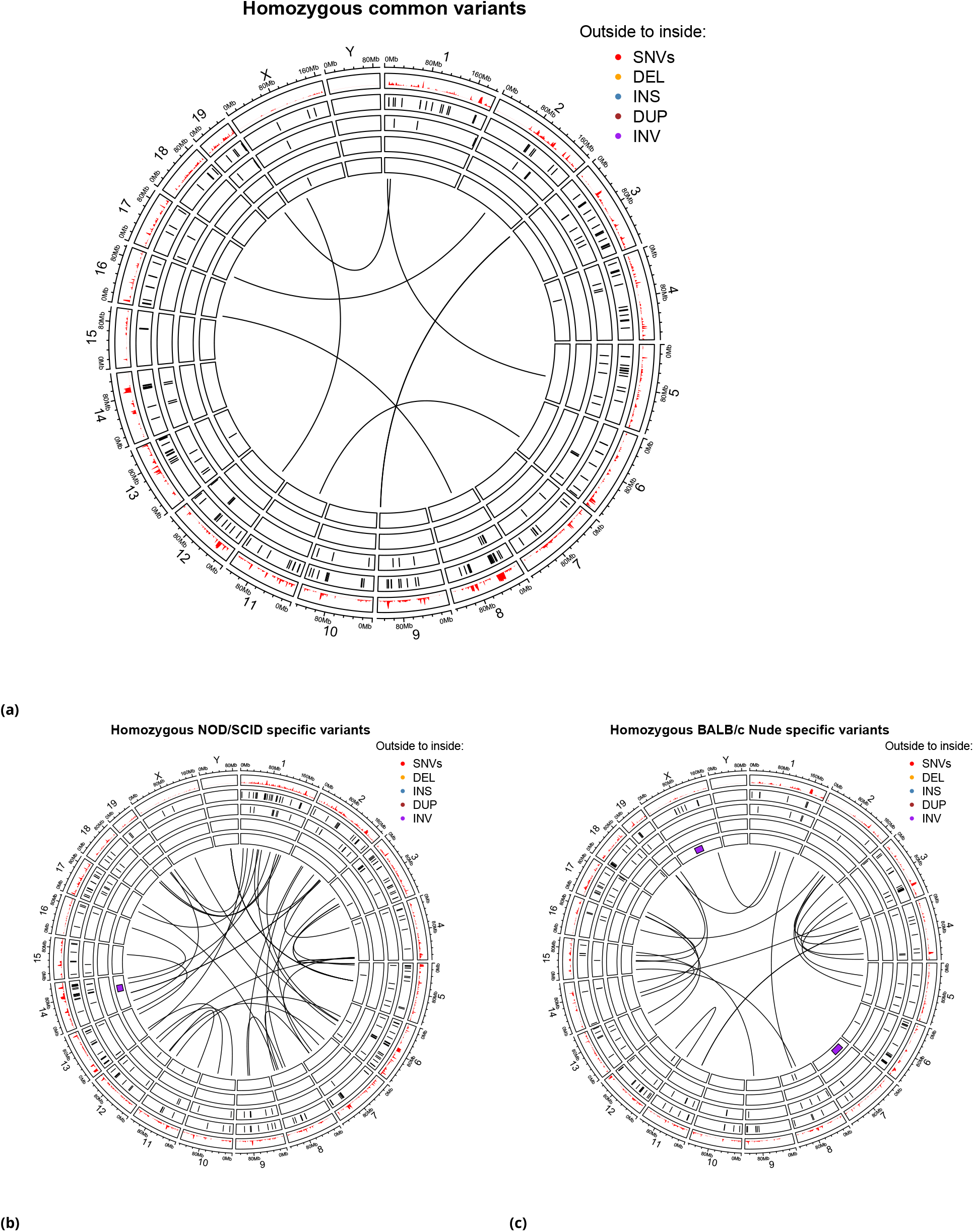
Systematic comparison of homozygous variants in BALB/c Nude and NOD/SCID mice. Homozygous SNVs and SVs were identified relative to the reference genome GRCm39 and were visualized in circular plots. **A)** Common variants in BALB/c Nude and NOD/SCID mice. Variants specific for NOD/SCID and BALB/c Nude strains were plotted in **B)** and **C)**, respectively. SNV density is reported in 1Mbp windows along the GRCm39 reference genome. Chromosomes are organized into sectors indicated by their label, start and end positions at the outside. Color codes indicate different types of variants (DEL=deletion,INS=insertion,DUP=duplication,INV=inversion in that order from the outer to the inner circle). Translocations are plotted in the middle of the circular plots as junctions between the breakpoints.

Few clusters can be noticed where a combination of apparently frequent events is occurring. For example, on chromosome 8 around 20 Mbp an accumulation of SNVs and SVs for both genotypes is noticeable (Figure 1a). One plausible reason for such accumulation of SNV calls in this region could be due to less accurate representation of the genomic architecture as supported by the nearby SV events. SNVs which are called in such regions might often be flawed through misrepresentation of the genomic structure and advocate for the need of strain specific genome assemblies. Only six translocations between chromosomes were detected to be common for both mouse strains in reference to the GRCm39 genome assembly (Figure 1a).

Analyzing the NOD/SCID specific variants (Figure 1b) we found a cluster of SNVs and SV on chromosome 7 around 100 Mbp. Interestingly, this region overlaps with the cluster of Olfr members (olfactory genes) which are known to be highly diversified in mice [***Young, 2002***]. Another example is a hot-spot of genomics rearrangements, containing a combination of inversion, deletion, and insertion events, that manifests itself on chromosome 14 around 80 Mbp. Such combination of different SV events often stem from the inability of the SV caller to resolve complex rearrangements and are better accessible through a *de novo* assembled genome instead.

### Strain specific genome assemblies

Based on the results of the genotyping, SV and SNV analysis, we pooled data from matching samples for each strain (Supplementary Table S1) and assembled the BALB/c Nude and NOD/SCID genomes. The coverage of non-human reads was equivalent to 44.5-fold and 47.2-fold the mouse genome size, respectively.

Due to the PDX nature of the samples it was important to verify the absence of human sequences pre-but as well post-assembly. The raw data were xengsorted and we aligned assembled contigs against the GRCm39 reference genome, screening non-mapping contigs further with the xengsort algorithm. Only contigs that could be clearly assigned as “host” genome-derived were kept. This subset was further screened against the RefSeq collection release 88 [***O’Leary et al., 2016***] using Mash Screen [***Ondov et al., 2019***], which did not identify any major contaminants such as e.g. bacterial genomes.

To further improve the continuity of the assembled HIFI contigs, we used Bionano optical maps as anchors for scaffolding. For each strain, new genomic DNA was prepared from a new PDX sample after confirming its genotype as described earlier. One Bionano flow-cell was used for each strain and raw molecules were competitively mapped against human and mouse *in silico* maps. Molecules with higher similarity to the host were then assembled and used for the scaffolding.

Bionano molecules of the NOD/SCID individual showed 15 % mouse-content and provided in total 185 Gbp of filtered molecules with a molecule N50 of 335 kbp. These molecules were assembled into 118 maps spanning 2.6 Gbp with a N50 of 74 Mbp. For the BALB/c Nude individual 75% of molecules were of mouse origin and they summed up in 770 Gbp filtered molecules with a N50 of 305 Kbp, which were assembled into 60 maps of 2.586 Gbp and a map N50 of 96.8 Mbp. These maps were then used to scaffold the HIFI assemblies into chromosome-scale sequences (Table 1).

**Table 1.** Basic genome assembly statistics. Statistics describing the genome assemblies for BALB/c Nude and NOD/SCID strains after cleaning and scaffolding with Bionano data.

| sample | scaffolds | unplaced contigs | Max. scaff.[Mbp] | Sum [Gbp] | NG50 [Mbp] |
| --- | --- | --- | --- | --- | --- |
| BALB/c Nude | 37 | 1033 | 167 | 2.75 | 167.3 |
| NOD/SCID | 28 | 270 | 195 | 2.72 | 121.8 |

The Bionano-based scaffolding resolved for both strains multiple chromosomes entirely, except for the telocentric centromeres and telomeres (Supplementary Figure S3 and Supplementary Figure S4). The unplaced contigs consist of difficult to resolve regions such as large repeats, mobile elements, and telomere/centromere-regions. The possibility that some haplotigs could be attributed to individual sample variation was not further investigated. The number of unplaced contigs was noticeably higher in the BALB/c Nude assembly compared to the NOD/SCID assembly (Table 1). We did not generate gene predictions but concentrated on lifting over known gene features to facilitate the adoption of the assemblies for the community. To that end, we used liftoff [***Shumate and Salzberg, 2021***] to find for each CDS of GRCm39 the boundaries matching on our assemblies and annotate them accordingly in a GTF file (Supplementary Table S2).

The BUSCO near-universal single-copy orthologs [***Manni et al., 2021a***] are often used to assess whether core genes are represented in an assembly and to what extent they are unique or repeated. The BUSCO results for the genome assemblies of both immunodeficient strains, BALB/c Nude and NOD/SCID, were on par with the results for the C57BL/6 reference (GRCm39) using either the genome assemblies (Table 2) or lift-over derived transcriptomes (Supplementary Table S3). BALB/c Nude had a slightly higher number of duplicated elements which might be contained within haplotigs that were not entirely purged.

**Table 2.** BUSCO scores for genome assemblies. BUSCO assessment of BALB/c Nude, NOD/SCID and C57BL/6 reference genome assemblies (GRCm39). Assemblies were analyzed against the glires_odb10 collection which contains 13’798 groups in total.

|  | BALB/c Nude | NOD/SCID | GRCm39 |
| --- | --- | --- | --- |
| Total Complete | 96.4 % | 96.5 % | 96.5 % |
| Complete single copy | 92.3 % | 93.6 % | 93.6 % |
| Complete duplicated | 4.1 % | 2.9 % | 2.9 % |
| Fragmented | 0.7 % | 0.6 % | 0.6 % |
| Missing | 2.9 % | 2.9 % | 2.9 % |

### Discordant regions between mouse strains

The gene annotation lift-over approach provides an estimate of the genes that are absent in our assemblies given the GRCm39 annotation. In total 981 and 1157 genes were missing for BALB/c Nude and NOD/SCID, respectively, after lifting over non-Y chromosome annotations from the reference genome. Out of these, 737 were common for both strains, most of which represented pseudogenes and long non-coding RNAs (see supplementary file “*common*_*missingLif tover*.*bed*”) as well as 89 protein-coding genes. Out of the missing protein-coding transcripts many are “Gm” prefixed gene names, indicating that they are potentially active transcripts which have no proper name for the moment. Excluding the Y chromosome genes, 98.1 % and 97.9% of GRCm39 annotated genes could be identified in BALB/c Nude and NOD/SCID assemblies, respectively.

Looking at the distribution of commonly missing genes along the chromosomes, one can identify clusters (Figure 2a). In regard to missing protein coding genes, chromosomes 7, 8 and 5 rank highest with 15,14 and 12 common missing features, respectively. Chromosome 7 is missing 6 members of the Scgb family (secretoglobin, family 1B), 3 Zscan members (Zinc finger and SCAN domain containing protein 4), 5 Vmn1r (vomeronasal 1 receptor) members and Gvin1 (GTPase, very large interferon inducible). Chromosome 8 misses for both strains almost exclusively Defa (defensin, alpha) members (Figure 2b) and Gm10131 (predicted pseudogene 10131). Finally, chromosome 5 is more heterogenous with 5 Pramel members (PRAME like), 4 Gm* and Vmn2r15 (vomeronasal 2, receptor 15), Cacna2d1 (Calcium Voltage-Gated Channel Auxiliary Subunit Alpha2delta 1) and AA792892 (expressed sequence AA792892).

**Figure 2.**
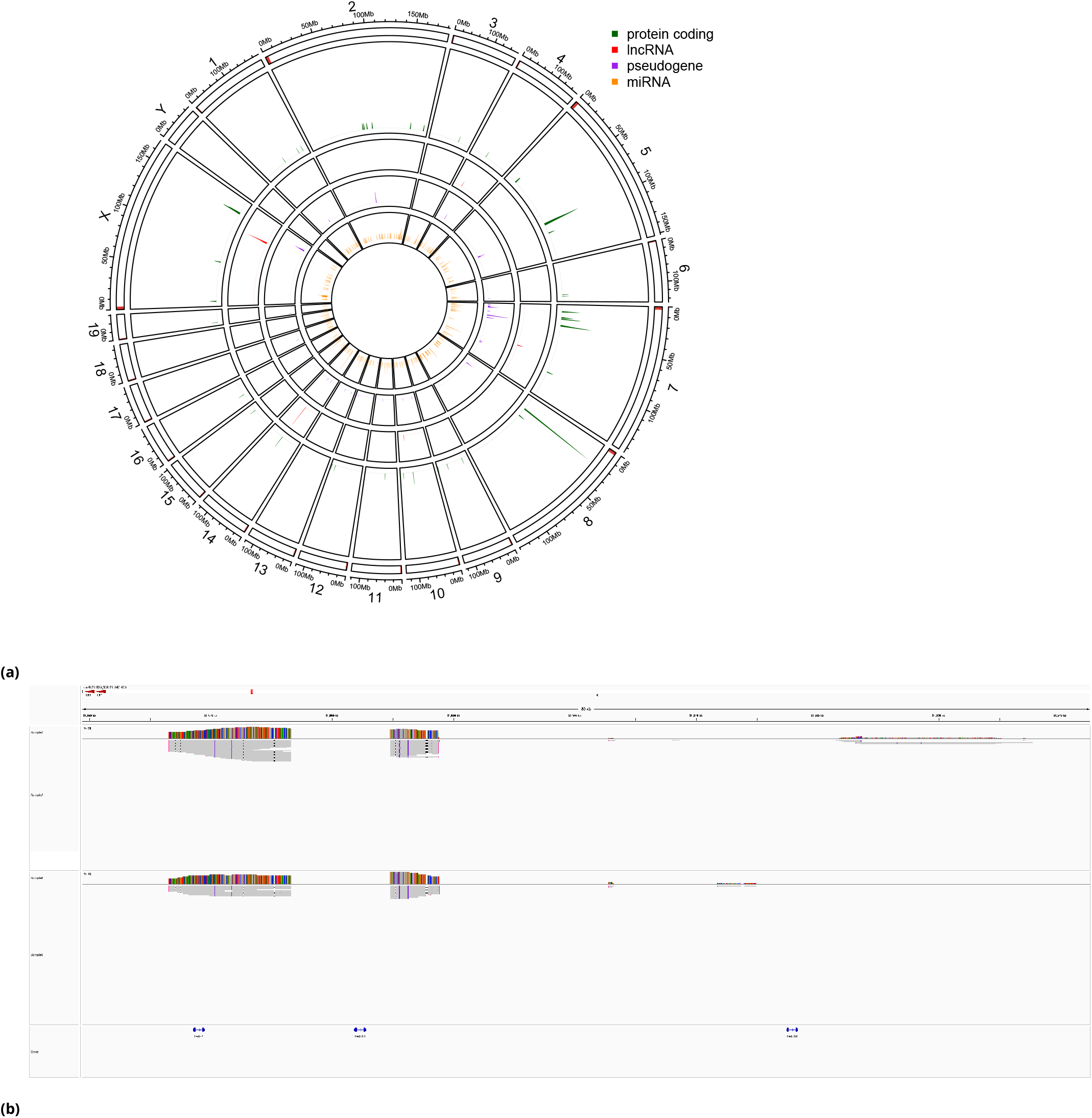
Identification of reference features that are absent in BALB/c Nude and NOD/SCID mice. **A)** Circular visualization of annotated reference features that could not be identified in the genome assemblies of BALB/c Nude and NOD/SCID strains. The organization of chromosomes is similar to that in Figure 1, except that Y chromosome was excluded and slices representing the five chromosomes with most number of missing protein coding features were enlarged for clarity. Color codes indicate types of features. **B)** Close-up view of the highlighted region containing members of the defensin alpha cluster on chromosome 8 (21,859,396-21,942,429; GRCm39). HIFI reads of one representative NOD/SCID and BALB/c Nude individual aligned to GRCm39 genome using minimap2 and visualized in IGV (Integrative Genomics Viewer) browser in upper and lower panels, respectively. Mismatch coloring was deactivated for clarity.

Looking at strain specific missing coding genes we found a BALB/c Nude specific absent Klra (known as well as Ly49) cluster on chromosome 6 around 130 Mbp (Figure 3a) which includes Klra10, Klra4, Klra6, Klra8 and Klra9. These genes are present in the NOD/SCID strain, but Klra homology compared to GRCm39 seems low, as indicated by many mismatches, and clipped long reads (Figure 3b). Composition of natural killer gene complex (NKC) genes and in particular Klar genes has been described to vary for different strains, with BALB/c Nude showing the most prominent differences and absence of Klra genes ***Higuchi et al. [2010***], which we can observe here as well.

**Figure 3.**
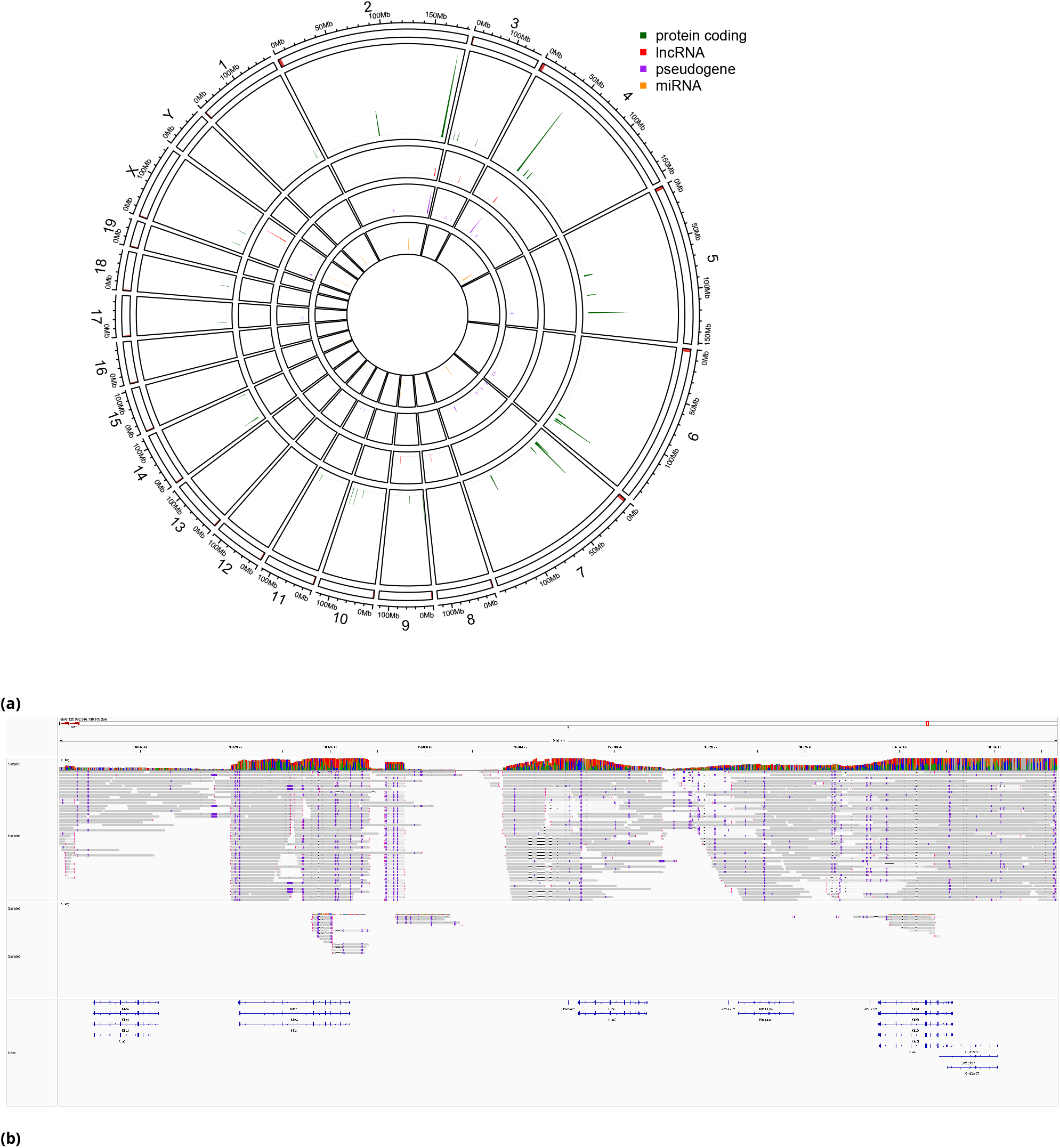
Identification of reference features that are specifically absent in BALB/c Nude strain. **A)** GRCm39 features that are absent specifically in BALB/c Nude genome assembly were plotted similarly to that in Figure 2a. B) Close-up view of Klra genes region on chromosome 6 (129,982,946-130,193,356; GRCm39). HIFI reads of two BALB/c Nude individuals were aligned to GRCm39 genome using minimap2 and visualized similarly to that in Figure 2.

## Conclusion

Here we assembled and annotated genomes for the NOD/SCID and BALB/c nude strains, characterizing most of the chromosomes to a chromosome-level scale. The vast majority of chromosomes are colinear with the existent C57BL/6 strain GRCm39 genome assembly. Both, raw-data SV analysis as well as genome comparison, identified though regions with large structural alterations, including olfactory genes and a BALB/c Nude specifically absent Klar gene cluster. The low homology Klar gene cluster in NOD/SCID, but especially the complete absence of multiple Klar genes of the NK cluster in BALB/c nude provide insights into the genomic alterations potentially explaining the previously observed reduced NK activity in this strain. The link between residual NK activity and reduced xentransplantation success remains the motivation of continuous constant effort to reduce NK activity further [***Panaampon et al., 2021***]. The availability of a genomic *de novo* assembly and thereby a genomic map provides the tools for a more profound studies of that immunological phenotype.

## Materials and Methods

### PacBio HIFI sequencing

High-molecular-weight gDNA was prepared from PDX tissue. The cells were lysed using Tissue Lyser II (Quiagen, Hilden Germany) and the DNeasy Blood & Tissue Kit 250 (Quiagen). The resulting gDNA was verified to be > 15 ug and > 10kbp long. DNA samples were purified and concentrated using AMPure PB beads (PacBio, Menlo Park, USA) to remove small fragments and impurities prior to library preparation. Library construction for HIFI protocol was done using SMRTbell Express Template Prep Kit 2.0 (PacBio), followed by another round of AMPure PB Beads Purification and Bluepippin size selection (< 8kbp). Samples were sequenced with Sequel II Sequencing Kit 2.0 on PacBio Sequel II (PacBio).

### Bionano sample and data preparation

DNA was extracted from tissue using the Bionano Prep SP Tissue and Tumor DNA Isolation Kit (Bionano, San Diego USA) and a Tissue Ruptor (Quiagen) targeting a final concentration of 50-120 ng/μL. DLE-1 labeling was done using the Bionano Prep DLS Labeling Kit with final concentration targeted at 4-12 ng/μL. Data collection was performed using Saphyr 2nd generation instruments.

### PacBio HIFI analysis

Consensus sequences were generated for all reads using ccs (v6.0.0; -j 0 –all –hifi-kinetics) and split into HIFI and low-quality reads using bamtools (***Barnett et al. [2011***]; v2.5.1; filter -tag “rq:<0.99”). uBAM entries were translated into fastq format using bam2fastq (v) and classified using xengsort (***Zentgraf and Rahmann [2021***]; v0.9.1; index generated using mouse genome and transcriptome as well as human genome and transcriptome : -p 4 –fill 0.94 -k 25 -P 4800000000 3FCVbb:u – hashfunctions linear945:linear9123641:linear349341847).

### Genotyping and SNV analysis

De novo variant calling was done using deepvariant (***Poplin et al. [2018***]; v1.1.0; model_type=PACBIO) using the HIFI host reads aligned to GRCm39 with pbmm2 (v1.4.0; –preset CCS). For the pileup-based analysis we used the tool abc (v0.2.1) providing the two marker mutation positions and the same alignments. For SNV analysis, regions with low complexity were removed using dust (v0.1; -w 64) and tandem repeat finder (v4.09; trf 2 5 7 80 10 50 2000 -f -d –m ; followed with TRFdat_to_bed.py) and bedtools (***Quinlan and Hall [2010***]; v2.29.0).

### SV analysis

Analysis was based on above-described alignments using pbsv (v2.4.0). Regions for which mapping quality was ≤ 5 for more than 2 reads within a range of 10bp were removed as well as SV smaller than 50bp, or > 5% allelic frequency or less than 3 reads supporting it (SURVIVOR; v1.0.7). Events in-between samples were compared using SURVIVOR merge within 1000bp range, same type and same direction of the event.

### Genome Assembly

The HIFI reads were assembled using hifiasm (***Cheng et al. [2021***]; yv0.12-r304) and were aligned against GRCm39 using mashmap (***Ondov et al. [2019***]; v2.0, -k 16 –s 1000 -f one-to-one –perc_identity 85). Contigs for which no alignment >= 5kbp was found were extracted (427 contigs in NOD/SCID mice and 1542 in BALB/c Nude) and screened again using xengsort to check if assembly generated contigs which now clearly identified as human derived. Contigs which were not clearly host assigned were removed. The remaining ones were screened using mash screen (v2.0, screen -w -p 30) against the refseq collection (RefSeq88n). It suggested there was a presence of endogenous mouse viruses in some, together with other murine viruses – but no graft similarities were anymore observed. These contigs were added back to the assemblies. We then used one round of haplotig purging as describe in purge_dups (v1.0.1).

### Bionano scaffolding

For the BALB/c Nude genotype the sample11 provided enough material to generate a matching Bionano run. For the NOD/SCID genotype this was not the case and we screened further 15 PDX for which lllumina WGS had been conducted. We identify a matching sample, Sample12, and run Bionano on it. We combined the genome sequence of the human and mouse genome and added in silico DLE-1 sites using their ‘ fa2cmap_multi_color.pl’ tool (Solve3.6.1_11162020). After de-noising we aligned the molecules against this combined reference and only kept molecules aligning preferentially to the mouse genome reference. These molecules were then assembled using Bionano’s tool ‘ pipelineCL.py’ (Solve3.6.1_11162020) and used to scaffold the matching, cleaned genome assembly. Assemblies are available under the BioProject PRJNA944543.

### Genome annotation

GRCm39 annotation (Mus_musculus.GRCm39.104.gtf_db) was lifted over using liftoff (v 1.5.1, -copies) for both strain genome assemblies.

### Genome assessment

BUSCO (***Manni et al. [2021b***]; v5.2.2) based on Glires dataset (containing 13798 Busco’s group on 24 genomes) was used to evaluate the completeness of each strain genome assembly. For the BUSCO transcriptome analysis lift over annotation were used together with gffread to extract the transcript FASTA entries (v0.12.7, feature “transcripts”). BUSCO transcript was run on these transcript catalogues (BALB/c Nude :140’147 transcripts, NOD/SCID : 139’795 transcripts, GRCm39: 142’434 transcripts).

## Author contributions statement

E.S.S. did the data-analysis. M.Q, H.T and B.C prepared PDX models, did tissue and DNA extraction as well as subsequent sequencing. E.S.S wrote the manuscript, B.A. and I.X. participated in writing. All authors reviewed the manuscript.

## Supplementary Material

**Table S1.** Characterization and genotyping of samples for genome assembly. Table of samples chosen for the genome assembly of BALB/c Nude and NOD/SCID genotypes. For each sample the mouse content and it’s genotype was analyzed based on WGS HIFI data. Genotype specific mutations in FOXn1 and prkdc genes were used for the genotyping.

| sample | Mouse-content | Genotype |
| --- | --- | --- |
| Sample1 | 25% | NOD/SCID |
| Sample2 | 22% | NOD/SCID |
| Sample3 | 62% | NOD/SCID |
| Sample4 | 32% | NOD/SCID |
| Sample5 | 31% | NOD/SCID |
| Sample6 | 33% | BALB/c Nude |
| Sample7 | 20% | BALB/c Nude |
| Sample8 | 26% | BALB/c Nude |
| Sample9 | 28% | BALB/c Nude |
| Sample10 | 19% | BALB/c Nude |
| Sample11 | 23% | BALB/c Nude |

**Table S2.** Liftover success of GRCm39 annotated gene-features for genome assembly of each strain. Liftover calculates gene coverage as the sum of all it’s transcripts covering the gene. From a total of 55416 genes in GRCm39, 1569 were removed prior lift-over as they were located on the Y-chromosome.

| strain | BALB/c Nude | NOD/SCID |
| --- | --- | --- |
| total genes | 53847 | 53847 |
| missing Genes | 981 | 1157 |
| lift-over genes | 52866 | 52690 |
| genes <90 % | 462 | 546 |
| genes >=90% & <95% | 201 | 237 |
| genes >=95% & <99% | 908 | 1023 |
| genes >=99% | 51295 | 50884 |

**Table S3.** BUSCO scores for annotation derived transcriptomes. BUSCO assessment of BALB/c Nude, NOD/SCID and C57BL/6 (GRCm39) reference transcriptomes. Transcriptome for BALB/c Nude and NOD/SCID were derived from lift-over annotation and all 3 were compared using “glires_odb10” collection which contains 13’798 groups in total.

|  | BALB/c Nude assembly | NOD/SCID assembly | GRCm39 |
| --- | --- | --- | --- |
| Total Complete | 99.7 % | 99.7 % | 99.9 % |
| Complete single copy | 39.8 % | 39.8 % | 39.9 % |
| Complete duplicated | 59.9 % | 59.9 % | 60 % |
| Fragmented | 0.1 % | 0 % | 0.0 % |
| Missing | 0.2 % | 0.3 % | 0.1 % |

**Figure S1.**
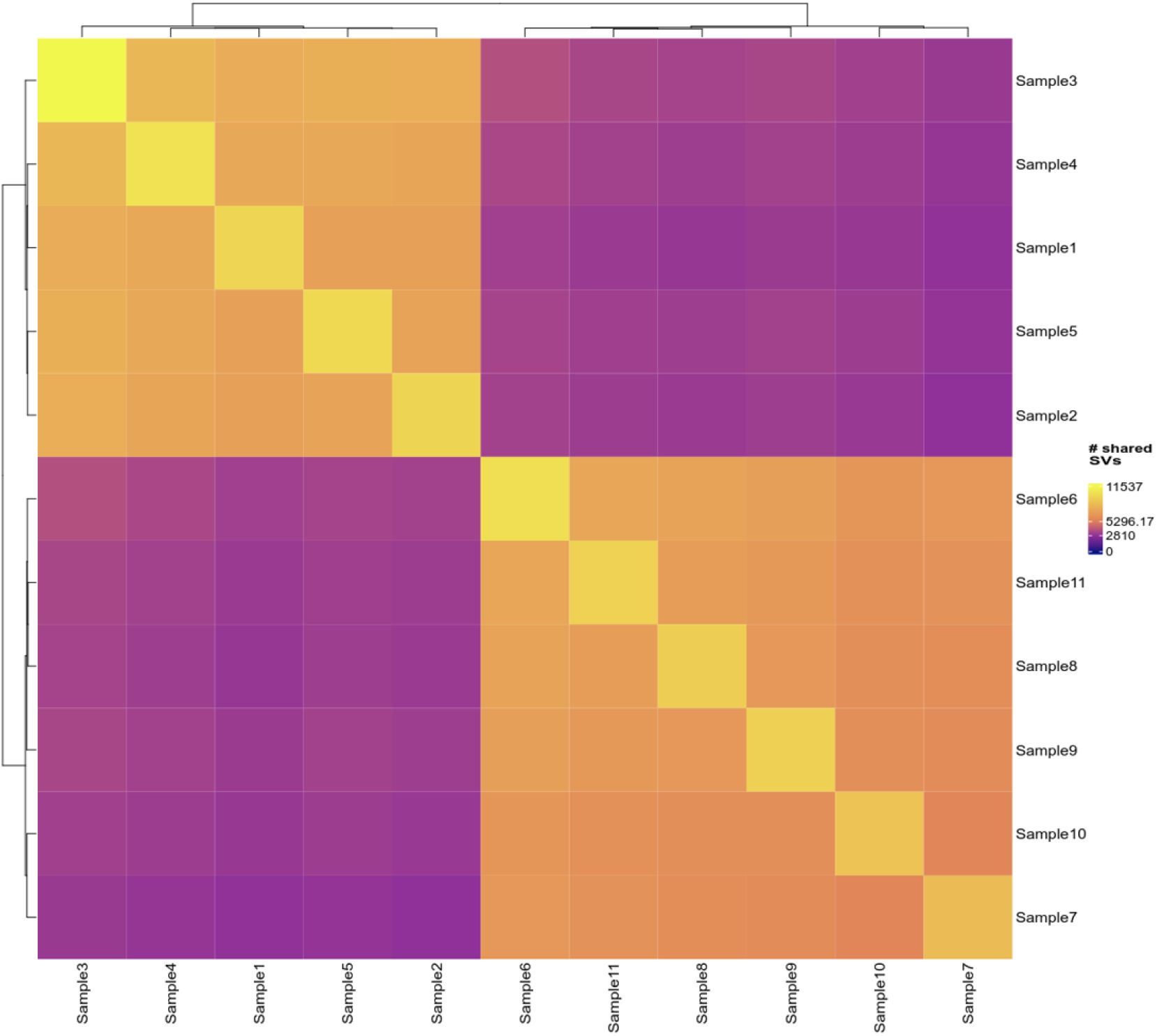
Structural variation events shared between samples. Heatmap comparing >1kbp homozygous SVs called individually for each sample using HIFI reads mapped against the GRCm39 genome assembly. Plotted are number of shared SVs between each sample. SVs are of same type, same direction and within a 1kbp maximal distance to each other.

**Figure S2.**
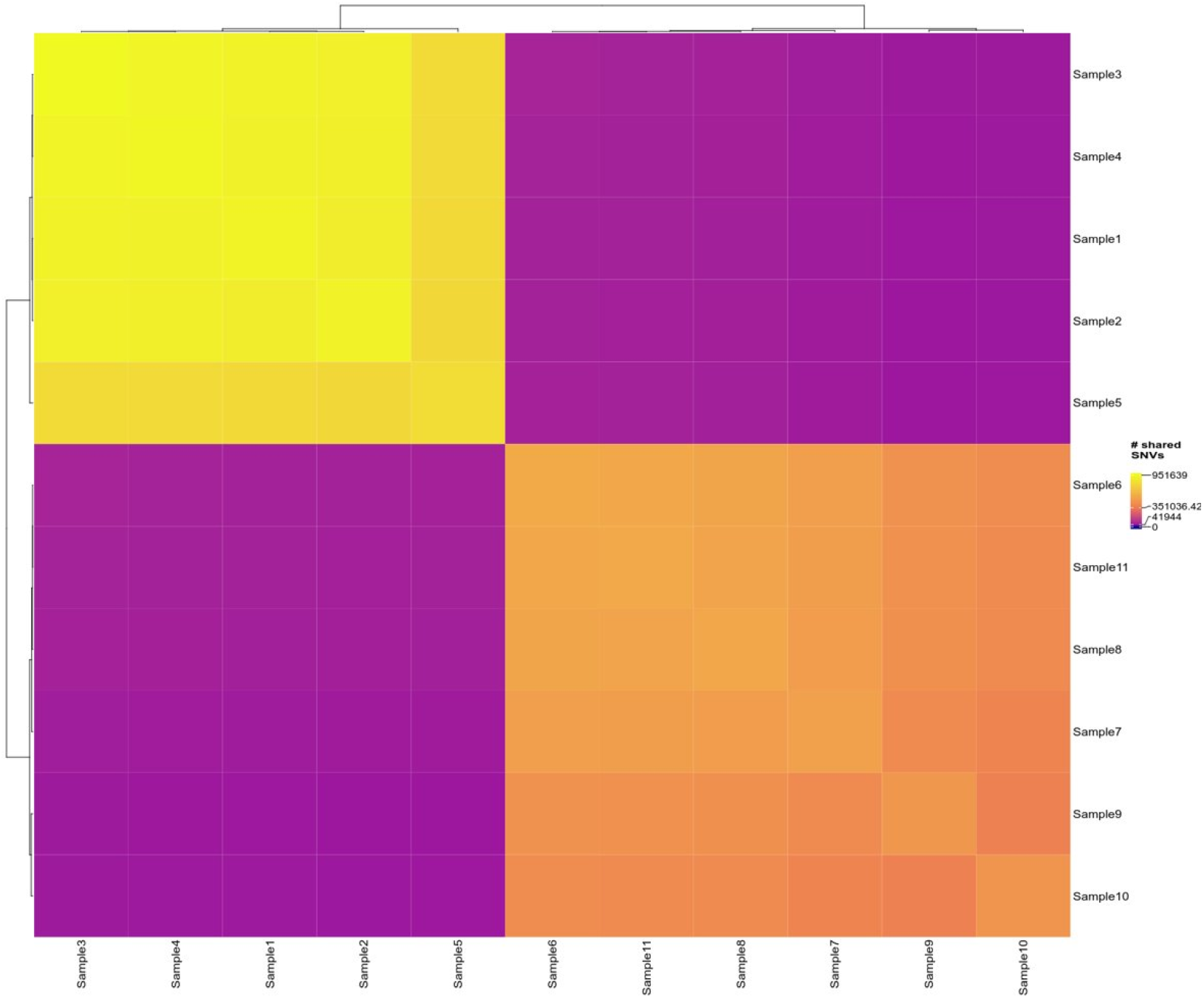
Single nucleotide variants shared between samples. Heatmap comparing homozygous SNVs between each sample using HIFI reads mapped against the GRCm39 genome assembly. Plotted are the number of shared SNVs between samples.

**Figure S3.**
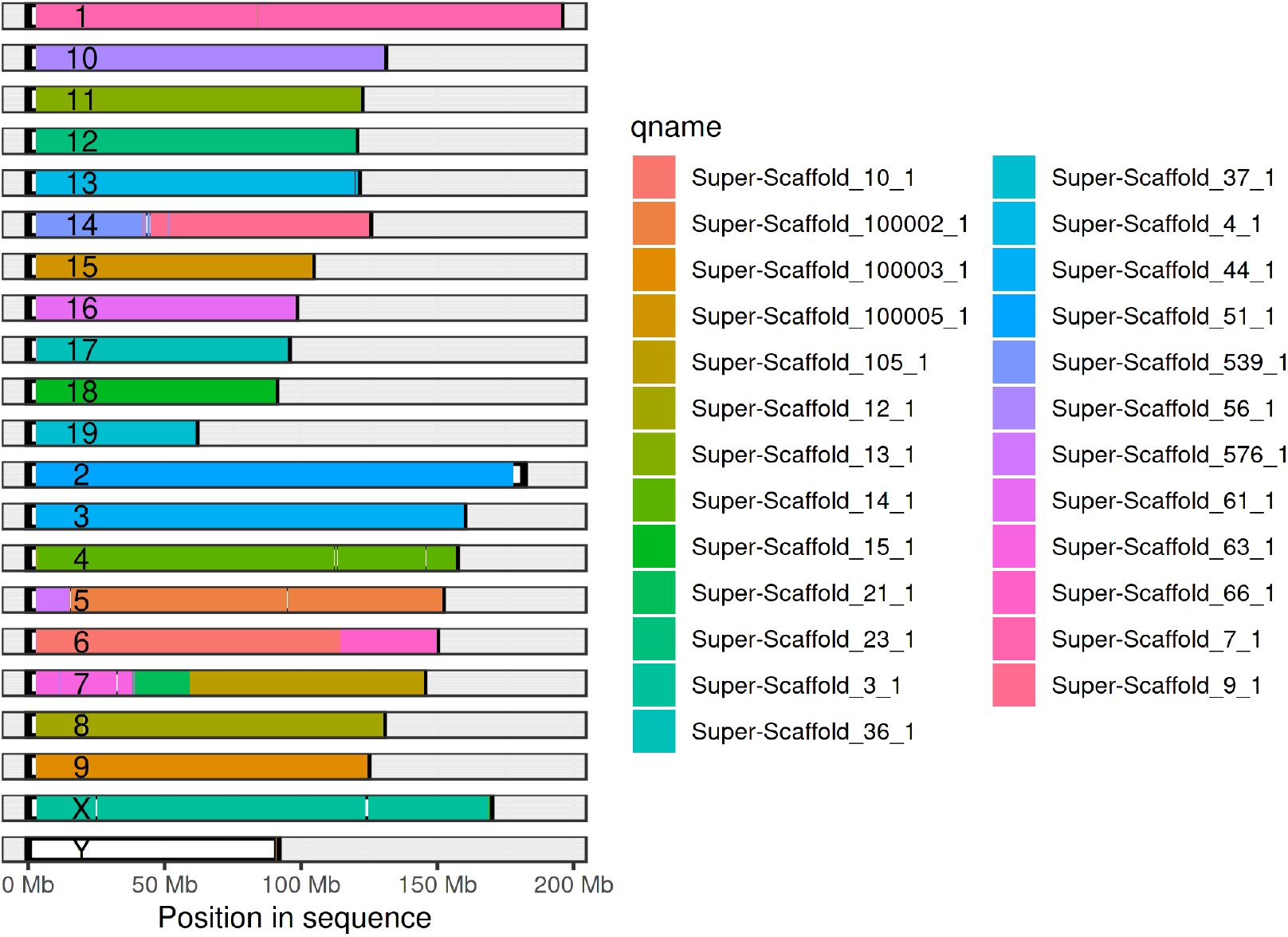
Coverage analysis of NOD/SCID genome assembly. Comparison of large segmental alignments between NOD/SCID genome assembly and GRCm39 reference assembly. Bold rectangles represent the GRCm39 chromosome space and color filling the matched assembly regions. Minimap2 alignment was done with “-x asm5”, alignments filtered for query and target contigs lengths > 1*e*7 and minimal match length > 1*e*4. Reference genome covered regions were visualized using pafr (version “115d2b1be6cdab5679a81a536200a03cef1ba0e9”).

**Figure S4.**
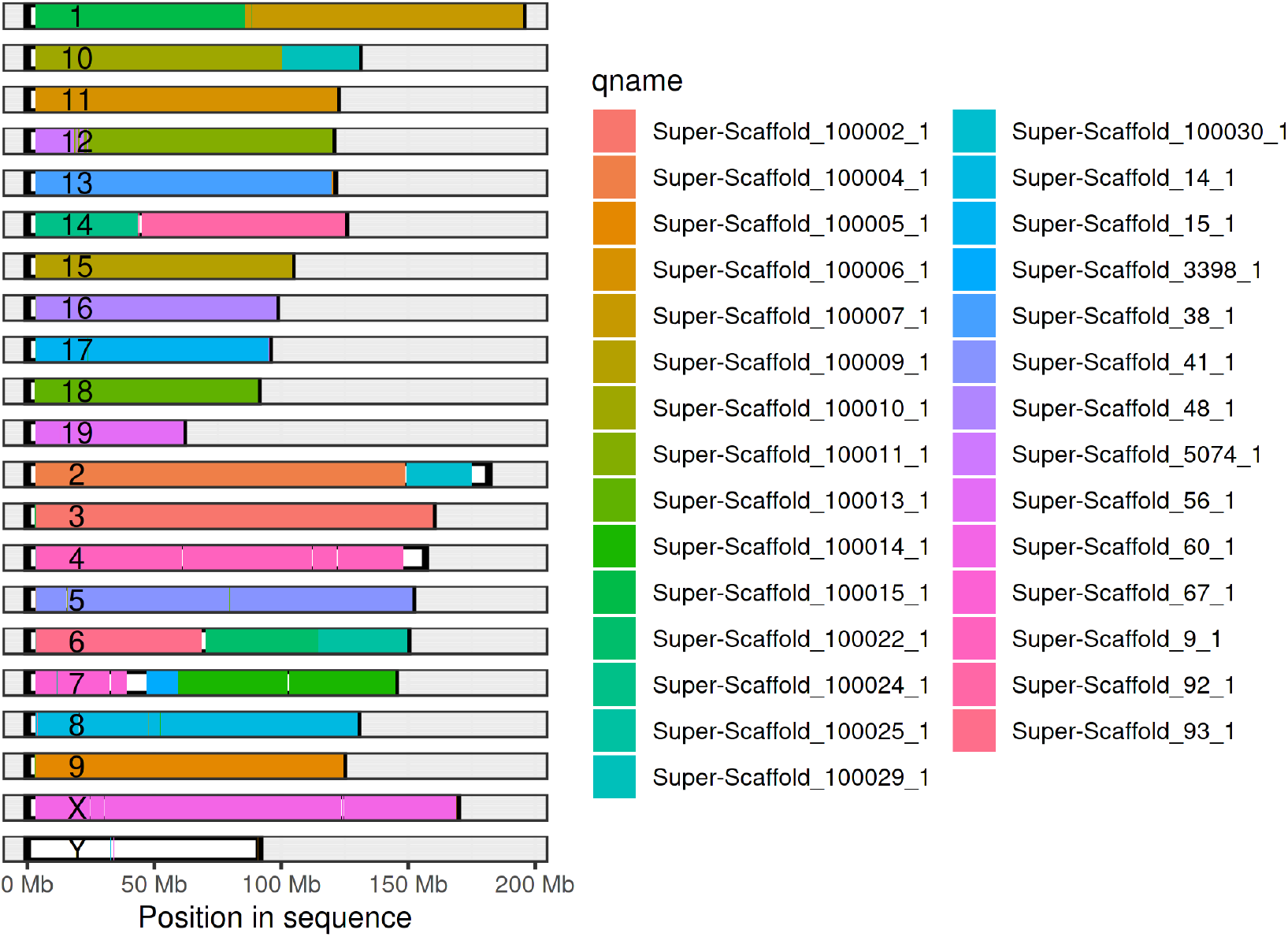
Coverage analysis of balb-c genome assembly. Comparison of large segmental alignments between balb-c genome assembly and GRCm39 reference assembly. Bold rectangles represent the GRCm39 chromosome space and color filling the matching assembly regions. Minimap2 alignment was done with “-x asm5”, alignments filtered for query and target contigs lengths > 1*e*7 and minimal match length > 1*e*4. Reference genome covered regions were visualized using pafr (version “115d2b1be6cdab5679a81a536200a03cef1ba0e9”).

## References

Barnett, D. W., Garrison, E. K., Quinlan, A. R., Strömberg, M. P., and Marth, G. T. (2011). BamTools: a C++ API and toolkit for analyzing and managing BAM files. Bioinformatics, 27(12):1691–1692.

Blunt, T., Gell, D., Fox, M., Taccioli, G. E., Lehmann, A. R., Jackson, S. P., and Jeggo, P. A. (1996). Identification of a nonsense mutation in the carboxyl-terminal region of DNA-dependent protein kinase catalytic subunit in the scid mouse. Proceedings of the National Academy of Sciences of the United States of America, 93(19):10285– 10290.

Bosma, G. C., Custer, R. P., and Bosma, M. J. (1983). A severe combined immunodeficiency mutation in the mouse. Nature, 301(5900):527–530.

Chen, C., Lin, W., Huang, Y., Chen, X., Wang, H., and Teng, L. (2021). The Essential Factors of Establishing Patient-derived Tumor Model. Journal of Cancer, 12(1):28–37.

Chen, X., Qian, W., Song, Z., Li, Q.-X., and Guo, S. (2020). Authentication, characterization and contamination detection of cell lines, xenografts and organoids by barcode deep NGS sequencing. NAR Genomics and Bioinformatics, 2(3):qaa060.

Cheng, H., Concepcion, G. T., Feng, X., Zhang, H., and Li, H. (2021). Haplotype-resolved de novo assembly using phased assembly graphs with hifiasm. Nature Methods, 18(2):170–175. Number: 2 Publisher: Nature Publishing Group.

Chuprin, J., Buettner, H., Seedhom, M. O., Greiner, D. L., Keck, J. G., Ishikawa, F., Shultz, L. D., and Brehm, M. A. (2023). Humanized mouse models for immuno-oncology research. Nature Reviews Clinical Oncology, pages 1–15. Publisher: Nature Publishing Group.

Ferraj, A., Audano, P. A., Balachandran, P., Czechanski, A., Flores, J. I., Radecki, A. A., Mosur, V., Gordon, D. S., Walawalkar, I. A., Eichler, E. E., Reinholdt, L. G., and Beck, C. R. (2022). Resolution of structural variation in diverse mouse genomes reveals chromatin remodeling due to transposable elements. Pages: 2022.09.26.509577 Section: New Results.

Gregory, S. G., Sekhon, M., Schein, J., Zhao, S., Osoegawa, K., Scott, C. E., Evans, R. S., Burridge, P. W., Cox, T. V., Fox, C. A., Hutton, R. D., Mullenger, I. R., Phillips, K. J., Smith, J., Stalker, J., Threadgold, G. J., Birney, E., Wylie, K., Chinwalla, A., Wallis, J., Hillier, L., Carter, J., Gaige, T., Jaeger, S., Kremitzki, C., Layman, D., Maas, J., McGrane, R., Mead, K., Walker, R., Jones, S., Smith, M., Asano, J., Bosdet, I., Chan, S., Chittaranjan, S., Chiu, R., Fjell, C., Fuhrmann, D., Girn, N., Gray, C., Guin, R., Hsiao, L., Krzywinski, M., Kutsche, R., Lee, S. S., Mathewson, C., McLeavy, C., Messervier, S., Ness, S., Pandoh, P., Prabhu, A.-L., Saeedi, P., Smailus, D., Spence, L., Stott, J., Taylor, S., Terpstra, W., Tsai, M., Vardy, J., Wye, N., Yang, G., Shatsman, S., Ayodeji, B., Geer, K., Tsegaye, G., Shvartsbeyn, A., Gebregeorgis, E., Krol, M., Russell, D., Overton, L., Malek, J. A., Holmes, M., Heaney, M., Shetty, J., Feldblyum, T., Nierman, W. C., Catanese, J. J., Hubbard, T., Waterston, R. H., Rogers, J., de Jong, P. J., Fraser, C. M., Marra, M., McPherson, J. D., and Bentley, D. R. (2002). A physical map of the mouse genome. Nature, 418(6899):743–750.

Higuchi, D. A., Cahan, P., Gao, J., Ferris, S. T., Poursine-Laurent, J., Graubert, T. A., and Yokoyama, W. M. (2010). Structural Variation of the Mouse Natural Killer Gene Complex. Genes and immunity, 11(8):637–648.

Karamboulas, C., Meens, J., and Ailles, L. (2020). Establishment and Use of Patient-Derived Xenograft Models for Drug Testing in Head and Neck Squamous Cell Carcinoma. STAR Protocols, 1(1):100024.

Li, Q.-X., Feuer, G., Ouyang, X., and An, X. (2017). Experimental animal modeling for immuno-oncology. Pharmacology & Therapeutics, 173:34–46.

Manni, M., Berkeley, M. R., Seppey, M., Simão, F. A., and Zdobnov, E. M. (2021a). BUSCO Update: Novel and Streamlined Workflows along with Broader and Deeper Phylogenetic Coverage for Scoring of Eukaryotic, Prokaryotic, and Viral Genomes. Molecular Biology and Evolution, 38(10):4647–4654.

Manni, M., Berkeley, M. R., Seppey, M., Simão, F. A., and Zdobnov, E. M. (2021b). BUSCO Update: Novel and Streamlined Workflows along with Broader and Deeper Phylogenetic Coverage for Scoring of Eukaryotic, Prokaryotic, and Viral Genomes. Molecular Biology and Evolution, 38(10):4647–4654.

Miga, K. H., Koren, S., Rhie, A., Vollger, M. R., Gershman, A., Bzikadze, A., Brooks, S., Howe, E., Porubsky, D., Logsdon, G. A., Schneider, V. A., Potapova, T., Wood, J., Chow, W., Armstrong, J., Fredrickson, J., Pak, E., Tigyi, K., Kremitzki, M., Markovic, C., Maduro, V., Dutra, A., Bouffard, G. G., Chang, A. M., Hansen, N. F., Wilfert, A. B., Thibaud-Nissen, F., Schmitt, A. D., Belton, J.-M., Selvaraj, S., Dennis, M. Y., Soto, D. C., Sahasrabudhe, R., Kaya, G., Quick, J., Loman, N. J., Holmes, N., Loose, M., Surti, U., Risques, R. a., Graves Lindsay, T. A., Fulton, R., Hall, I., Paten, B., Howe, K., Timp, W., Young, A., Mullikin, J. C., Pevzner, P. A., Gerton, J. L., Sullivan, B. A., Eichler, E. E., and Phillippy, A. M. (2020). Telomere-to-telomere assembly of a complete human X chromosome. Nature, 585(7823):79–84. Number: 7823 Publisher: Nature Publishing Group.

Okada, S., Vaeteewoottacharn, K., and Kariya, R. (2019). Application of Highly Immunocompromised Mice for the Establishment of Patient-Derived Xenograft (PDX) Models. Cells, 8(8):889.

O’Leary, N. A., Wright, M. W., Brister, J. R., Ciufo, S., Haddad, D., McVeigh, R., Rajput, B., Robbertse, B., Smith-White, B., Ako-Adjei, D., Astashyn, A., Badretdin, A., Bao, Y., Blinkova, O., Brover, V., Chetvernin, V., Choi, J., Cox, E., Ermolaeva, O., Farrell, C. M., Goldfarb, T., Gupta, T., Haft, D., Hatcher, E., Hlavina, W., Joardar, V. S., Kodali, V. K., Li, W., Maglott, D., Masterson, P., McGarvey, K. M., Murphy, M. R., O’Neill, K., Pujar, S., Rangwala, S. H., Rausch, D., Riddick, L. D., Schoch, C., Shkeda, A., Storz, S. S., Sun, H., Thibaud-Nissen, F., Tolstoy, I., Tully, R. E., Vatsan, A. R., Wallin, C., Webb, D., Wu, W., Landrum, M. J., Kimchi, A., Tatusova, T., DiCuccio, M., Kitts, P., Murphy, T. D., and Pruitt, K. D. (2016). Reference sequence (RefSeq) database at NCBI: current status, taxonomic expansion, and functional annotation. Nucleic Acids Research, 44(D1):D733–D745.

Olson, B., Li, Y., Lin, Y., Liu, E. T., and Patnaik, A. (2018). Mouse Models for Cancer Immunotherapy Research. Cancer Discovery, 8(11):1358–1365.

Ondov, B. D., Starrett, G. J., Sappington, A., Kostic, A., Koren, S., Buck, C. B., and Phillippy, A. M. (2019). Mash Screen: high-throughput sequence containment estimation for genome discovery. Genome Biology, 20(1):232.

Panaampon, J., Sasamoto, K., Kariya, R., and Okada, S. (2021). Establishment of Nude Mice Lacking NK Cells and Their Application for Human Tumor Xenografts. Asian Pacific Journal of Cancer Prevention : APJCP, 22(4):1069– 1074.

Poplin, R., Chang, P.-C., Alexander, D., Schwartz, S., Colthurst, T., Ku, A., Newburger, D., Dijamco, J., Nguyen, N., Afshar, P. T., Gross, S. S., Dorfman, L., McLean, C. Y., and DePristo, M. A. (2018). A universal SNP and small-indel variant caller using deep neural networks. Nature Biotechnology, 36(10):983–987. Number: 10 Publisher: Nature Publishing Group.

Quinlan, A. R. and Hall, I. M. (2010). BEDTools: a flexible suite of utilities for comparing genomic features. Bioinformatics, 26(6):841–842.

Rossello, F. J., Tothill, R. W., Britt, K., Marini, K. D., Falzon, J., Thomas, D. M., Peacock, C. D., Marchionni, L., Li, J., Bennett, S., Tantoso, E., Brown, T., Chan, P., Martelotto, L. G., and Watkins, D. N. (2013). Next-Generation Sequence Analysis of Cancer Xenograft Models. PLOS ONE, 8(9):e74432. Publisher: Public Library of Science.

Schlake, T. (2001). The nude gene and the skin. Experimental Dermatology, 10(5):293–304. _eprint: https://onlinelibrary.wiley.com/doi/pdf/10.1034/j.1600-0625.2001.100501.x.

Shumate, A. and Salzberg, S. L. (2021). Liftoff: accurate mapping of gene annotations. Bioinformatics, 37(12):1639–1643. Publisher: Oxford Academic.

Wenger, A. M., Peluso, P., Rowell, W. J., Chang, P.-C., Hall, R. J., Concepcion, G. T., Ebler, J., Fungtammasan, A., Kolesnikov, A., Olson, N. D., Töpfer, A., Alonge, M., Mahmoud, M., Qian, Y., Chin, C.-S., Phillippy, A. M., Schatz, M. C., Myers, G., DePristo, M. A., Ruan, J., Marschall, T., Sedlazeck, F. J., Zook, J. M., Li, H., Koren, S., Carroll, A., Rank, D. R., and Hunkapiller, M. W. (2019). Accurate circular consensus long-read sequencing improves variant detection and assembly of a human genome. Nature Biotechnology, 37(10):1155–1162. Number: 10 Publisher: Nature Publishing Group.

Woo, X. Y., Srivastava, A., Graber, J. H., Yadav, V., Sarsani, V. K., Simons, A., Beane, G., Grubb, S., Ananda, G., Liu, R., Stafford, G., Chuang, J. H., Airhart, S. D., Karuturi, R. K. M., George, J., and Bult, C. J. (2019). Genomic data analysis workflows for tumors from patient-derived xenografts (PDXs): challenges and guidelines. BMC Medical Genomics, 12(1):92.

Young, J. M. (2002). Different evolutionary processes shaped the mouse and human olfactory receptor gene families. Human Molecular Genetics, 11(5):535–546.

Zentgraf, J. and Rahmann, S. (2021). Fast lightweight accurate xenograft sorting. Algorithms for Molecular Biology, 16(1):2.

